# Differential detection of *Entamoeba* species in stool samples collected from children in District Swat, Khyber Pakhtunkhwa Pakistan

**DOI:** 10.1101/729798

**Authors:** Muhammad Iftikhar Khan, Sumaira Shams, Asar Khan, Ali Akbar, Ijaz Muhammad, Atta Ullah, Muhammad Inam, Abid Ali

## Abstract

**Background:** Amoebiasis is an intestinal disease caused by enteric protozoan called *Entamoeba histolytica* belongs to the Genus *Entamoeba*. The main reason of infection is the contamination of food and water due to the poor sanitation. Among *Entamoeba* species*, Entamoeba histolytica* is highly pathogenic while the other species are non-pathogenic and needs no medical treatment.

**Methodology:** A total of 400 stool samples were collected from different areas of district Swat and were processed for screening of amoebic cells. Microscopically identified samples containing amoebic cells were stored at −20 °C till DNA extraction. Extracted DNA was used in a PCR reaction with specific reference primers to amplify the target DNA.

**Results:** Out of all 400 stool samples 111 (27.7%) were found positive through microscopy while PCR reaction confirmed 80 out of microscope positive samples. Among 80 PCR positive samples, the infection with *Entamoeba dispar* was most common (57.5%) followed by *E. histolytica (*47.5%) and *Entamoeba moshkovskii* (20%). The positive cases for mono-infection of *E. dispar* were 33 (41.25%), followed by *E.histolytica* 25 (31.25%) and *E. moshkovskii* 7 (8.75%). The co-infection of *E. histolytica* with *E. dispar* and *E. moshkovskii* was 6 (7.5%) and 2 (2.5%), respectively. Similarly the co-infection of *Entamoeba dispar* with *Entamoeba moshkovskii* was also 2 (2.5%) while 5 (6.25%) samples were observed with mixed infection of *E. histolytica, E. dispar* and *E. moshkovskii*.

**Significance of the study:** The aim of the study was to detect and differentiate the *E. histolytica, Entamoeba dispar* and *Entamoeba moshkovskii* using conventional microscopy and polymerase chain reaction. The results suggested that the use of PCR is necessary to differentiate *E. histolytica* from *E. dispar* and *E. moshkovskii* and therefore, to avoid unnecessary treatment the present study recommend the use of PCR for the routine diagnosis of amoebiasis in the study area. It is also suggested that further studies from this area may also facilitate the understanding of genetic diversity of these pathogens.

## Introduction

*Entamoeba histolytica* is an enteric protozoan belongs to genus *Entamoeba* which is the causative agent of amoebiasis [1]. The Genus *Entamoeba* comprises of several species among them six reside in human intestinal tract including *E. histolytica, E. dispar, E. moshkovskii, E. coli, E. poleki* and *E. hartmanni* [2]. There are two forms *Entamoeba,* the trophozoite which has short life span and are mobile that can invade the different organ systems, while the cysts are long surviving form of *Entamoeba* that colonize the patient. The cyst consists of resistant walls which protect these species from desiccation in external environment [3]. In external environment the cysts can survive for several days, weeks or month mostly in damp conditions and are responsible for transmission of infection [4].

The transmission of cyst occurs with the ingestion of contaminated food or water and trophozoite proliferate in the lumen of large intestine that may cause disease [5]. Mostly These luminal parasite live commensally and feed on bacteria but eventually it cause injury to the epithelial layer of large intestine and can reach the organs such as liver, brain and lungs by ulcerating the mucosal tissues [6]. The major symptoms of amoebiasis includes diarrhea, amoebic dysentery and liver abscess [7]. The amoebic infections may be symptomatic or asymptomatic and can cause a lot of clinical manifestation but majority of infected individual are asymptomatic [8]. The size of cyst ranges from 10-15 μm in diameter while the size of trophozoit is 10-60 μm in diameter. The cyst in mature form release from large intestine in feces and can remain viable for several days in cool and moist environment, in water the cyst can survive for up to 30 days and die when the temperature is below 5 °C [9].

There are 34 to 50 million symptomatic cases reported every year worldwide, which result in 40 to 100 thousands deaths annually [10]. The amoebic infection is common in humans and non-human primates and consider as the third leading cause of death after malaria and schistosomiasis [11]. The prevalence of amoebiasis varies from 4-81% worldwide and the estimated data shows that 10% of the people are infected with amoebic infections globally. Previous studies have shown that this infection is prevalent in Asia, Africa, South and Central America [12]. The poor hygienic conditions and environmental pollution have been shown to strengthen amoebic infections [13]. Mostly the amoebic cases are reported from developing countries and are introduced to developed countries due to the moments of human and animals [14]. The factors like poverty, pollutions, over population, poor education, unhygienic food and unsanitary conditions facilitate the transmission of disease. It is contagious and can spread from person to person that enforce accurate and appropriate diagnosing, treatment and prevention [15].

The diagnosis of amoebiasis faces challenge and most of the individuals receive the unnecessary treatment with anti-amoebic drugs. Morphologically the *E. histolytica* is indistinguishable from *E. dispar* and *E. movshkovskii* therfore conventional microscopic method cannot differentiate them [16]. Light microscope is mostly used to confirm amoebiasis by staining and wet smear but the results mostly mislead to differentiate the trophozoite and cyst of pathogenic *E. histolytica* from the cyst and trophozoite of non-pathogenic *E. dispar* and *E. moshkovskii* which infect humans occasionally [17]. But this traditional parasitological method have the advantage that they are less costly and not require the expensive chemicals and equipment’s. If a trained microscopist is available these analysis can be performed easily [18]. Now a days there are many methods use for diagnosing amoebiasis like iso-enzyme assay, rapid indirect haemagglutination, assay (IHA), culture using Dr. Bohlav biphasic and Boek amoebic medium [19]. Other diagnostic methods like culturing, ELISA and PCR are used for diagnosing purposes. Serological tests can diagnose *E. histolytica* infections but it is not used in routine laboratory based identification because due to time consuming (takes up to 12 weeks). Now a days there is no commercial and accurate molecular methods available other than PCR that can distinguish *E. histolytica*, *E. moshkovskii* and *E. dispar* [20].

To prevent the amebiasis at present the interruption of the fecal-oral spreading of infectious cyst stage of the parasite is necessary. As the cysts are resistant to low doses of iodine and chlorine, in developing countries it is necessary to boil the water before drinking.

The present study was designed to avoid excessive and unnecessary treatment with anti-protozoal drugs for *Entamoeba* species infected individuals, and to provide better understanding of the epidemiology of these parasites in the human population. Studies on human protozoal infections are very limited from Pakistan especially from rural populations. Therefore, it is necessary to identify and differentiate the pathogenic *Entamoeba* species and to evaluate the prevalence of these parasites in children of district Swat, Khyber Pakhtunkhwa (KP), Pakistan through conventional microscopic and advance molecular techniques.

## Research methodology

### Study area description

District Swat KP is a river valley and 165 kilometer away from the capital city, Peshawar. There are several mountain peaks ranging from 4500 to 6000 meter above the sea level. The elevation of Swat valley at south is over 600 meter high and rapidly rises toward the north side. The temperature of this region ranges from −2-33 °C. This area is predominantly rural and mostly the residents live in small villages [21]. The present study was carried out in District Swat, KP to investigate the differential detection of *Entamoeba* species in stool samples of children. The stool samples were collected from labs and randomly from the people of small villages and towns like Odigram, Kanju, Saidu Sharif and Naway Kali/Mingora (Fig. 1).

**Fig. 1.**
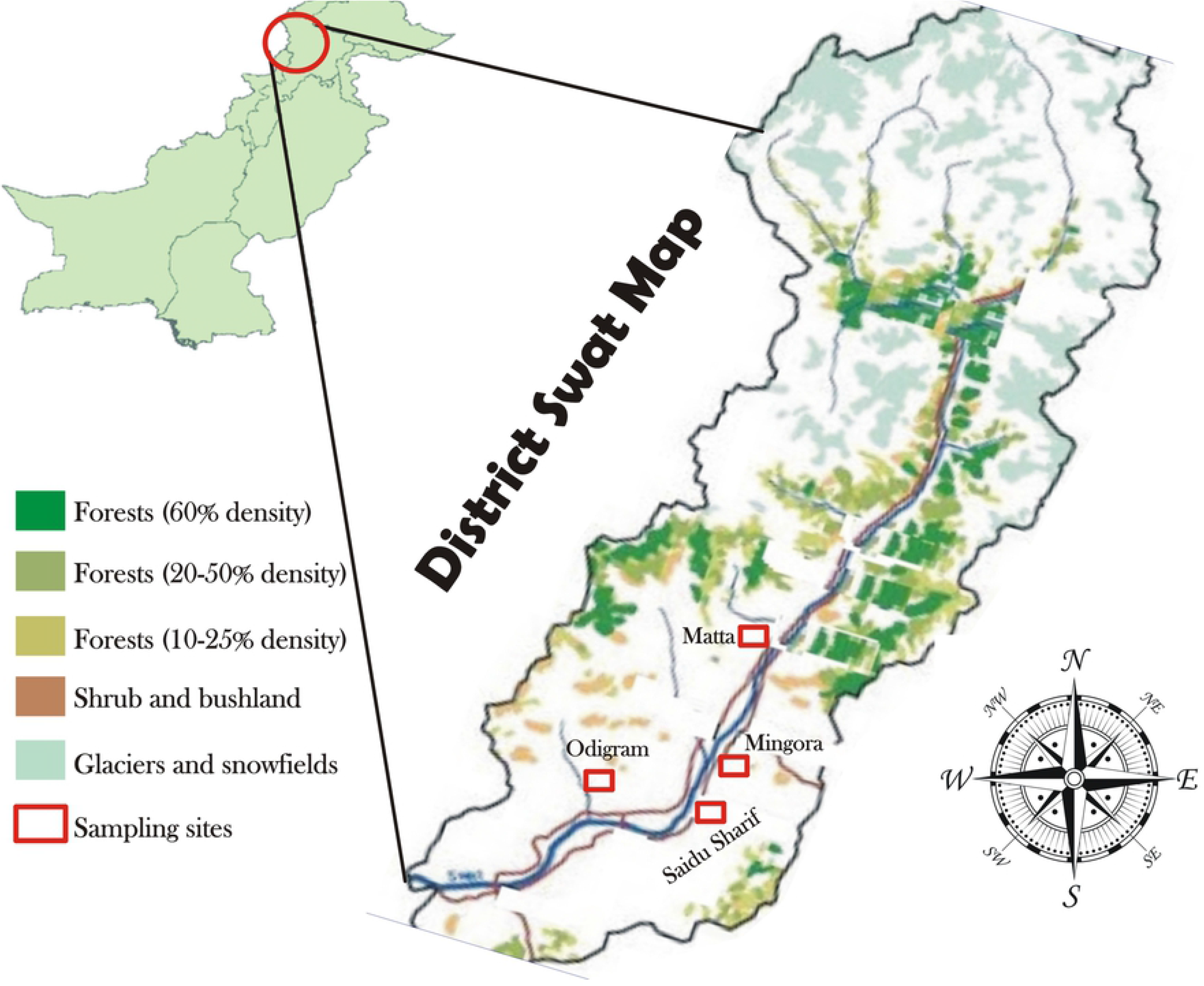
Map of district Swat (draw and modified by the authors using Coral Draw 9.0 and Photoshop 7.0 software’s) showing the study area and sampling sites (highlighted).

### Ethical approval of the study

The ethical approval for study design and the written consent procedures was obtained from Research Ethics Committee (REC) Department of Zoology Abdul Wali Khan University Mardan. Permission was obtained from the heads of Hospitals, Health care centers and families of the study participants. To facilitate the sampling from patients attending the hospitals and health care centers, the families of all participants were also informed verbally about the purpose of the study. All the responses were recorded using a standard questionnaire designed for this study.

### Sample collection

A total of 400 stool samples were collected from children (Age 3-12 years) with the assistance of their parents by visiting different areas and labs of District Swat. For the detection of *Entamoeba* species the samples were kept in 5 ml falcon tubes containing 2% potassium dichromate and 70% ethanol as a preservative. The sample containers were labeled (name, age, sex, area and month) and were processed to the Parasitology laboratory of Zoology Department, Abdul Wali Khan University Mardan.

### Laboratory analysis of samples

#### Iodine Wet Mount

A total of 2 mg stool samples were mixed with Lughole’s iodine. After mixing a drop of sample was placed on glass slide and the cover slip was placed over the suspension for microscopic analysis [22].

#### Staining procedure

A single drop (20-30 μl) of stool sample was placed on a clean slide and was left to air dry for 10-15 minutes. After drying the smear a drop of methanol was poured on the slide for fixation. A drop of Giemsa stain was added on dried slide and was left for 5 minutes and finally the slide was washed with distilled water and placed under microscope for examination.

#### Morphological identification

After preparation of iodine wet mount for laboratory observations each slide was further examined and confirmed under microscope using 10 X and 40 X lenses for the presence or absence of *Entamoeba*. Fecal samples that containing amebic cells were photographed using a digital camera (SONY, Japan) and were stored at −20 ºC until DNA extraction.

#### DNA extraction and amplification of PCR product

The genomic DNA was extracted using, GF-1 DNA extraction kit (Vivantis Technologies, Malaysia). The given standard protocol was used for DNA extraction with minor modifications. The reference primers of specific gene (16S rRNA) were used to amplify the purified DNA (Table 1).

**Table 1.**
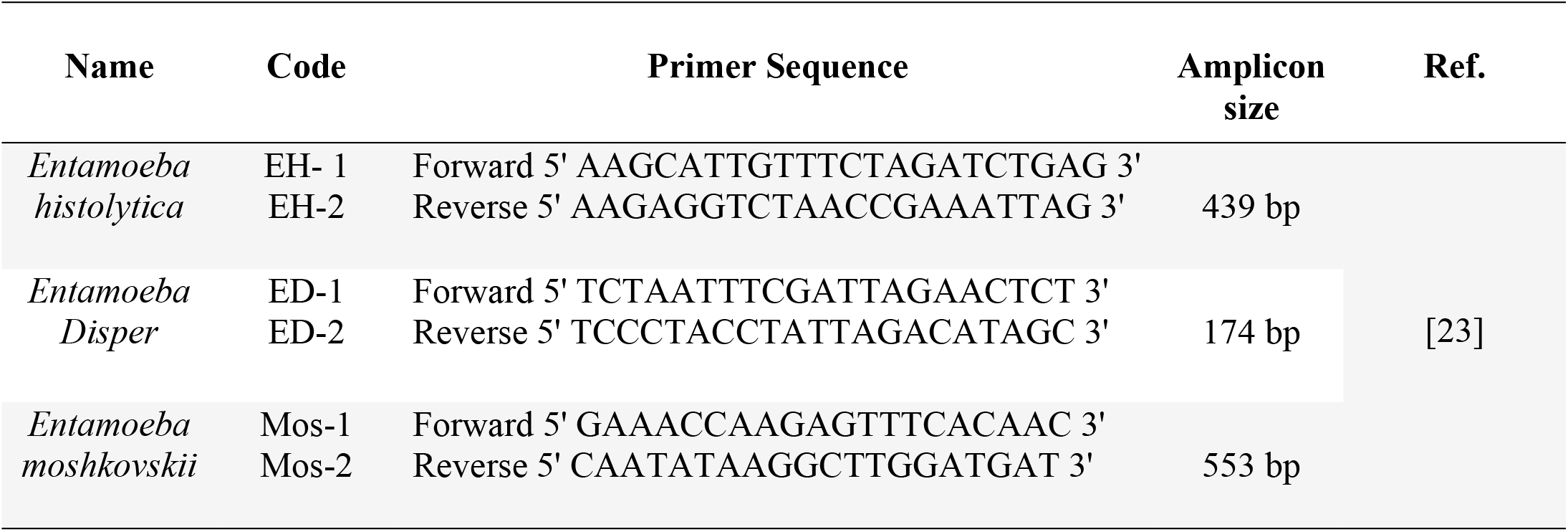
The reference primers set used for *Entameoba* specie specific amplification.

The PCR amplification reaction was performed using a thermal cycler (BIO-RAD T100, USA). The amplification was done as described previously with minor modifications [24]. A 0.3 μl *Taq* DNA polymerase was added with 5 μl extracted DNA, 2.2 μl *Taq* Buffer, 2.4 μl MgCl_2_ (25mM), 1.0 μl dNTPs and 1.0 μl of each forward and reverse primers. Amplification conditions consisted of 1 cycle of 3 minutes at 94 °C, 30 cycles of 30 seconds at 94 °C, 30 seconds at 55 °C and 30 seconds at 72°C and the final stage was consisted of 1 cycle of 7 minutes at 72°C. The PCR products were then subjected to gel electrophoresis in 2% agarose, staining with ethidium bromide and visualization in a UV trans-illuminator (Bio-Doc, California, USA). The DNA amplified fragments of each sample was determined by identifying the 439 bp band for *E. histolytica,* 174 bp for *E. dispar* and 553 bp for *E. moshkovskii* compare with 50 bp DNA ladder (Fermentas Germany), used as size maker.

#### Statistical data management and graphical analysis

All the statistical data collected in the field were keyed in MS-Excel sheet to identify inconsistencies in the data, consistency checks were done e.g. whether dates of children were less than 2 and greater than 5 years. Duplicate checks were performed and if found, these were removed. The data was exported to Graphpad Prism V.7.0 (CA, USA) for data cleaning by running frequencies for the different variables. A statistical significant level of 5% was used with a 95% confidence interval (C.I). Continuous variables were summarized using the mean. The prevalence was calculated as a proportion of children who tested positive for *Enatameoba* species. The odd ratios and *p-values* were used to determine whether each of the included variable has an effect on the prevalence of *Enatameoba*. While, the graphical representation, cropping and editing of figures (map) was done through Adobe Photoshop 7.0 (USA) and Coral Draw 9.0 (SYNEX, Brooklyn, NY).

## Results

Out of all 400 stool samples, 110 were collected from Odigram which showed 27.2% (no. 30) prevalence of *Entamoeba*. Similarly 110 out of 400 stool samples were collected from Naway Kali/Mingora in which the prevalence of *Entamoeba* was 30.90% (no. 34 cases) while 96 out of 400 stool samples were collected from Saidu Sharif showed 28.12% (27) prevalence of *Entamoeba* and 84 stool samples collected from Kanju showed 23.80% (in 20 cases) (Table 2).

**Table 2.** Overall observed prevalence (%) of *Entamoeba* species in selected sites of the study area

| Name of Area | Total samples examined | Observed positive | Prevalence (%) |
| --- | --- | --- | --- |
| Odigram | 110 | 30 | 27.2 % |
| Naway kali/Mingora | 110 | 34 | 30.90 % |
| Saidu sharif | 96 | 27 | 28.12 % |
| Kanju | 84 | 20 | 23.80 % |
| Sum | **400** | **111** | **27.75%** |

Among the total examined stool samples, 111 (27.7 %) samples showed the cyst or trophozoit of any of the *E. histolytica, E. disper* or *E. moshkovskii* species (Fig 2).

**Fig 2.**
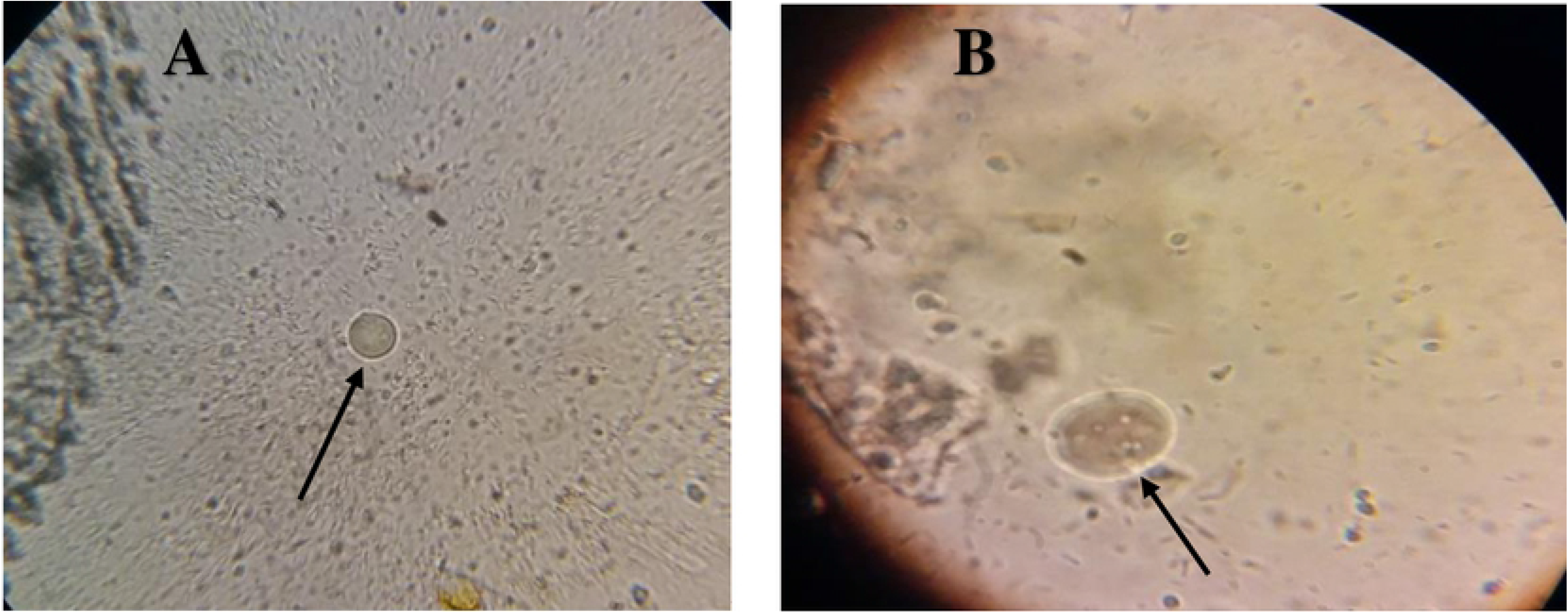
Microscopic detection and identification of *Entameoba* spp. (A) 10x magnification (B) 40x magnification power lenses.

### Gender wise prevalence of *Entamoeba* species

A total of 210 out of 400 stool samples were collected from male children from different areas of Swat among which the prevalence of *Entamoeba* species were observed 33.33 % (70/210) and 190 out of 400 samples were collected from female in which the prevalence of *Entamoeba* was 21.57 % (36/190). A 110 samples were collected from Odigram among which the prevalence of *Entamoeba* in male was 33.84 % (22/65) and in female the prevalence was 24.44% (11/45) while 110 stool samples were collected from Naway Kali/Mingora among which the prevalence of *Entamoeba* in male was 31.74% (20/63) and in female the prevalence was 25.53% (12/47). Similarly 96 stool samples were collected from Saidu Sharif in which 36.17% (17/47) in male and 22.44% (11/49) prevalence of *Entamoeba* was recorded in female while in Kanju out of 84 stool samples the 31.42% (11/35) in male and 14.28% (7/49) in female were positive for *Entamoeba* spp. (Table 3).

**Table 3.**
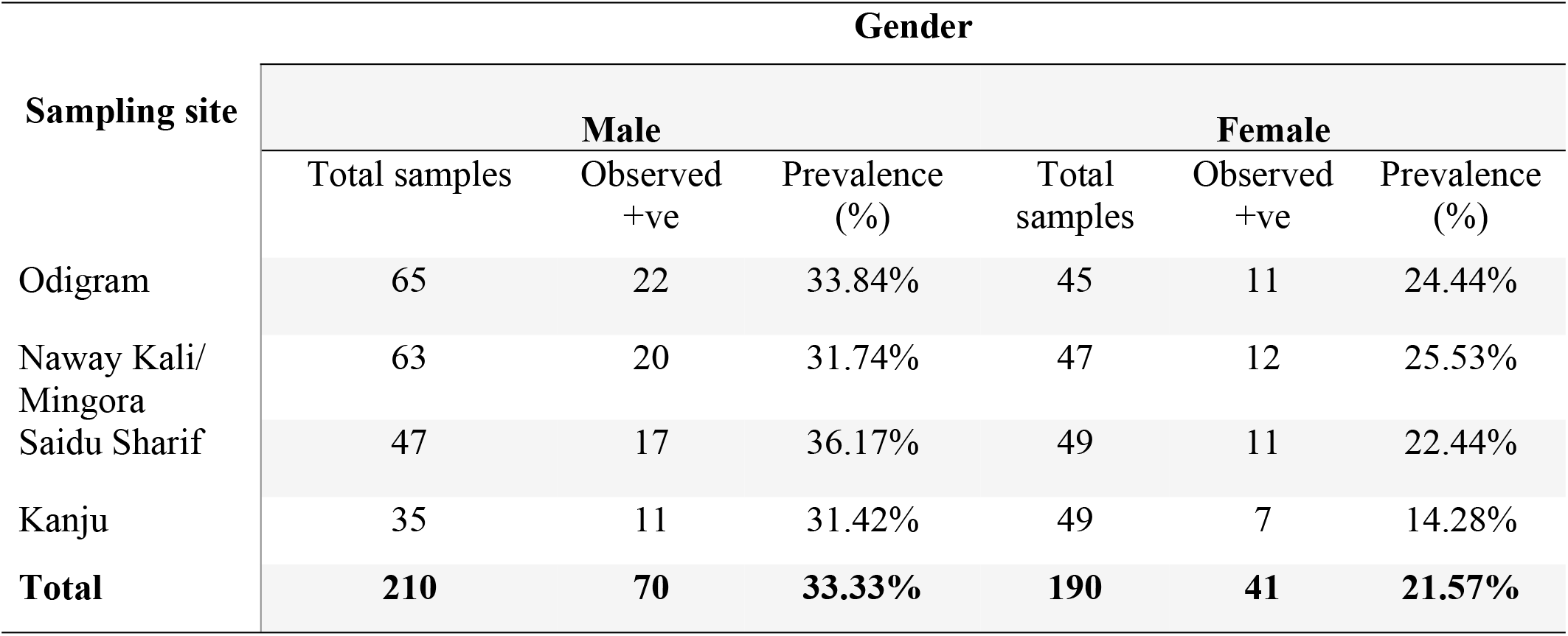
Gender wise observed prevalence (%) of *Entamoeba* spp. in the study area (n = 400).

| Sampling site | Gender |  |  |  |  |  |
| --- | --- | --- | --- | --- | --- | --- |
|  | Male |  |  | Female |  |  |
|  | Total samples | Observed +ve | Prevalence (%) | Total samples | Observed +ve | Prevalence (%) |
| Odigram | 65 | 22 | 33.84% | 45 | 11 | 24.44% |
| Naway Kali/<br>Mingora | 63 | 20 | 31.74% | 47 | 12 | 25.53% |
| Saidu Sharif | 47 | 17 | 36.17% | 49 | 11 | 22.44% |
| Kanju | 35 | 11 | 31.42% | 49 | 7 | 14.28% |
| **Total** | **210** | **70** | **33.33%** | **190** | **41** | **21.57%** |

In the gender wise prevalence, the male population (Mean= 52.5, SD= ±17.5) was observed to be susceptible to the infection with a high prevalence of 33.33% (70/210) and a significant *p-value* of 0.01. Female (Mean= 47.5, SD= ±10.25) infected cases revealed no statistical significance (*p-value* =0.08) correlation with prevalence rate.

### Prevalence of *Entamoeba* complex in different age groups

The children were divided into two groups, the one group was aged below 5 years while the second group included children of 5 to 12 years of age. A total of 170 out of 400 stool samples were collected from children aged below 5 which showed the prevalence of *Entamoeba* complex 22.35 % (38/170) and 230 samples were collected from children aged 5-12 among them the prevalence of *Entamoeba* was 31.73 % (73/230). Out of all 400 samples 110 were collected from Odigram which showed the prevalence of *Entamoeba* 34.48% (20/58) in children aged 5-12 and 19.23% (10/52) in children aged below 5 years (Table 4).

**Table 4.**
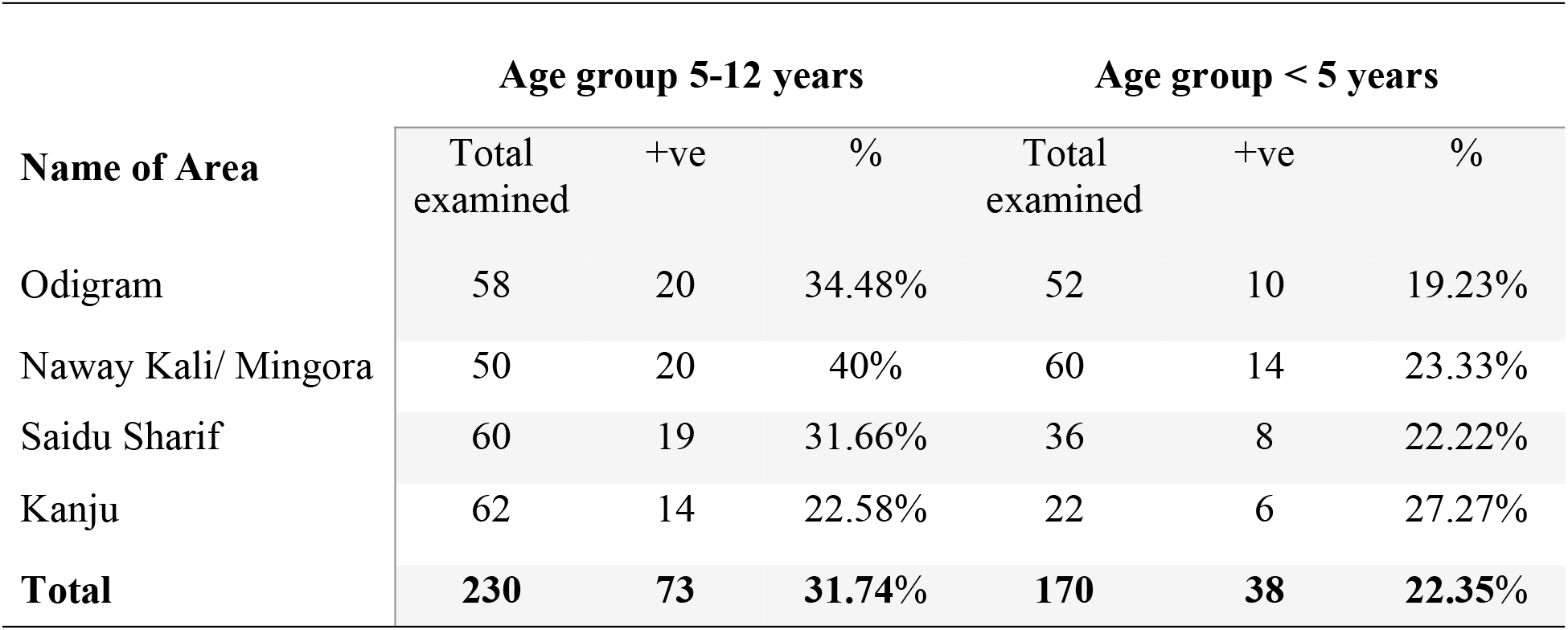
The distribution of *Entameoba* species among different age groups.

| Name of Area | Age group 5-12 years |  |  | Age group < 5 years |  |  |
| --- | --- | --- | --- | --- | --- | --- |
|  | Total examined | +ve | % | Total examined | +ve | % |
| Odigram | 58 | 20 | 34.48% | 52 | 10 | 19.23% |
| Naway Kali/ Mingora | 50 | 20 | 40% | 60 | 14 | 23.33% |
| Saidu Sharif | 60 | 19 | 31.66% | 36 | 8 | 22.22% |
| Kanju | 62 | 14 | 22.58% | 22 | 6 | 27.27% |
| **Total** | **230** | **73** | **31.74%** | **170** | **38** | **22.35%** |

Similarly 110 samples were collected from Naway Kali/Mingora in which the prevalence of *Entamoeba* complex in children aged 5-12 years was 40% (20/50) and in children below 5 years age was 23.33% (14/60). A total of 96 samples from Saidu Sharif showed the prevalence of *Entamoeba* species in 31.66% (19/60) cases in age group 5-12 years and 22.22% (8/36) prevalence was observed in children aged below 5 years 27 (28.12%). A total of 84 stool samples were collected from Kanju among which 22.58% (14/62) were observed positive for *Entamoeba* in age group 5-12 while the prevalence of *Entamoeba* in children aged below 5 years was recorded 27.27% (6/22). Statistically, no correlation observed in the age groups < 5 years of age (Mean= 18.25, SD= ±2.87) with a *p-value* of 0.096. While a high statistical variation was observed in age group 5-12 years of age (Mean= 9.5, SD= ±3.42) with a 0.001 of *p* value.

### Seasonal dynamics of *Entamoeba* species

The prevalence of infection was 12.66% (10/79) in March among total 79 collected. Out of 79 samples collected in April, the prevalence of infection was 16 (20.25%). Similarly a total of 80 samples were collected in May in which 23 (28.75%) were positive for *Entamoeba* while in June 30 (37.04%) out of 81 were positive for *Entamoeba* species. Out of 81 samples collected in the month of July, 32 (39.51%) were positive for *Entamoeba* species through microscopy (Fig 3).

**Fig 3.**
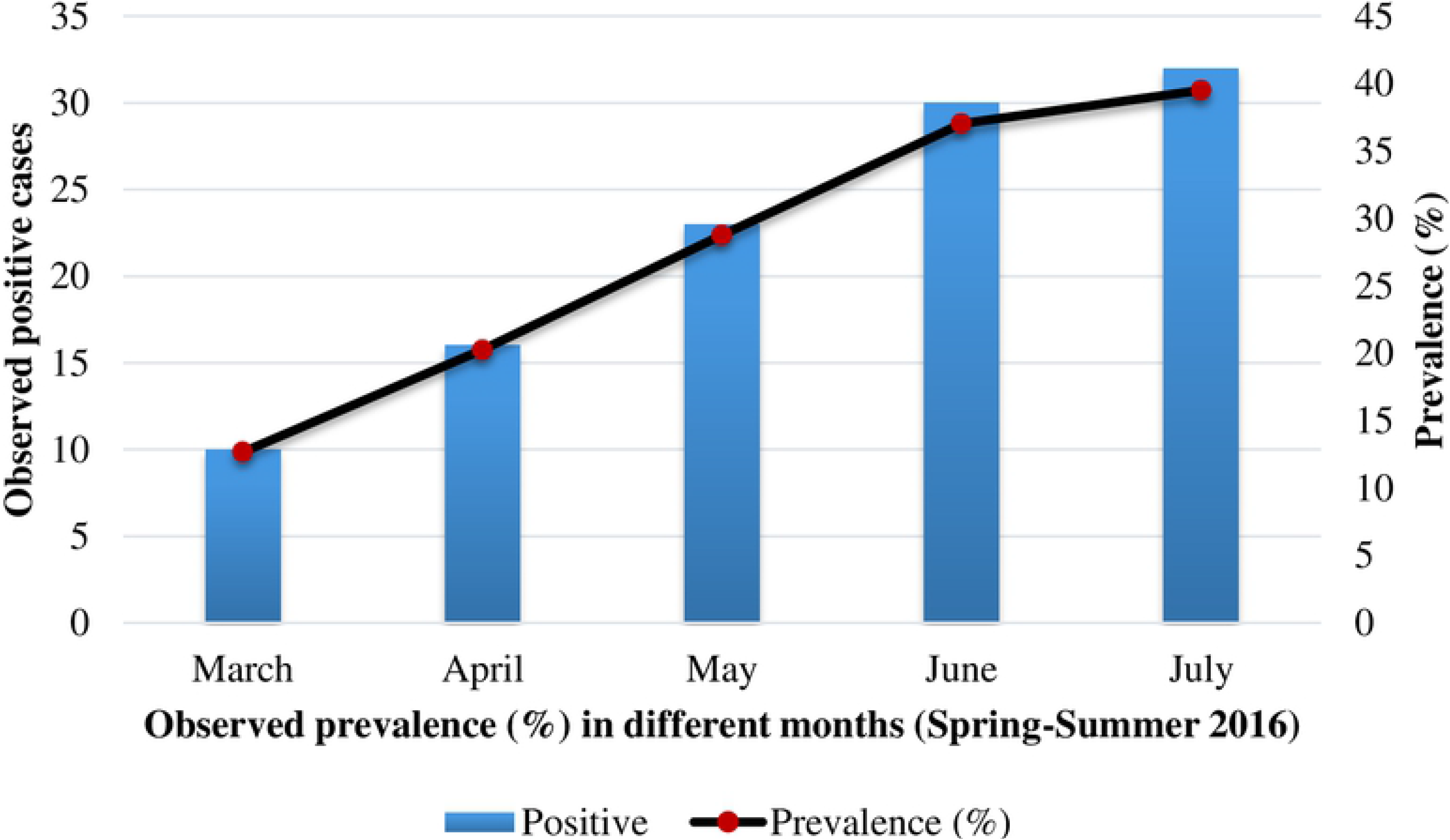
Observed prevalence (%) in different months (from Spring 2016 to summer 2016)

### Genotypic confirmation and prevalence (%) of *Entamoeba* species

All samples observed positive (111/400) through microscopy were subjected to PCR for the confirmation and identification of *Entamoeba* species. It was confirmed that only 80 of the microscopic positive samples were successfully amplified by PCR assay. Among PCR positive samples, 31.25 % (25/80) were positive for *E. histolytica,* 41.25% (33/80) were positive for *E. disper* while 8.75% (7/80) were positive for *E. moshkovskii*. The mixed infection of *E. histolytica* with *E. disper* or with *E. moshkovskii* and *E. dispar* with *E. moshkovskii* were also observed. The PCR-based confirmed co-infection of *E. histolytica* with *E. disper* and *E. moshkovskii* was 6.25% (5/80) (Table 5).

**Table 5.**
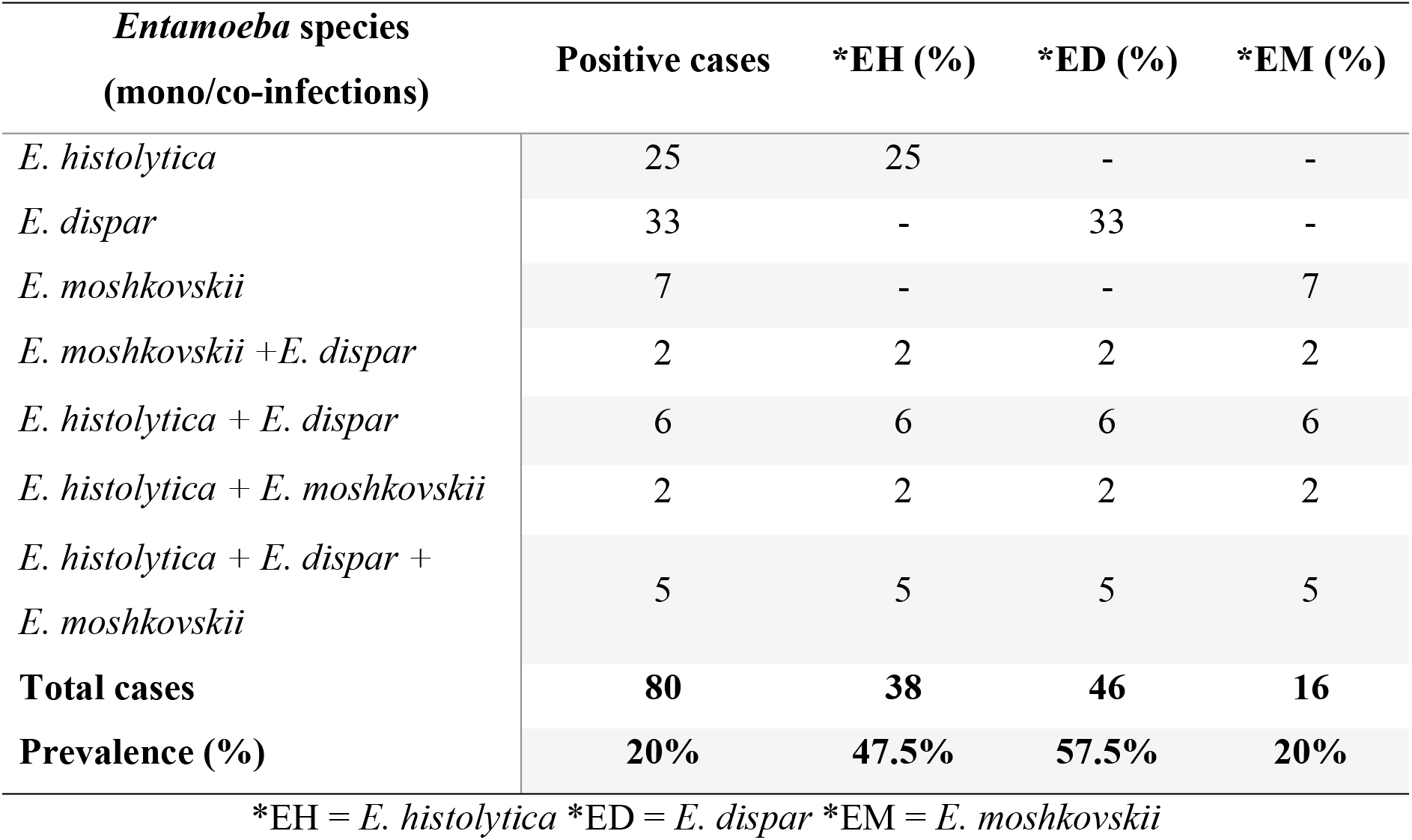
PCR based prevalence and mono/co-infection of *E.histolytica, E.dispar* and *E. moshkovskii*.

The coinfection of *E. histolytica* with *E. dispar* was 7.5% (6/80) while the coinfection of *E. histolytica* with *E. moshkovskii* was 2.5% (2/80) and the coinfection of *E. dispar* with *E. moshkovskii* was also 2.5% (2/80). Overall *E. histolytica* positive samples including single and coinfection were 47.5% (38/80) and *E. dispar* positive samples including coinfection were 57.5% (46/80) while the *E. moshkovskii* monoinfection was recorded 20% (16/80). All PCR based positive samples for cysts/trophozoite of *E*.either for *E. histolytica, E. dispar* or *E. moshkovskii* that were also positive by microscopy (Fig. 4).

**Fig 4.**
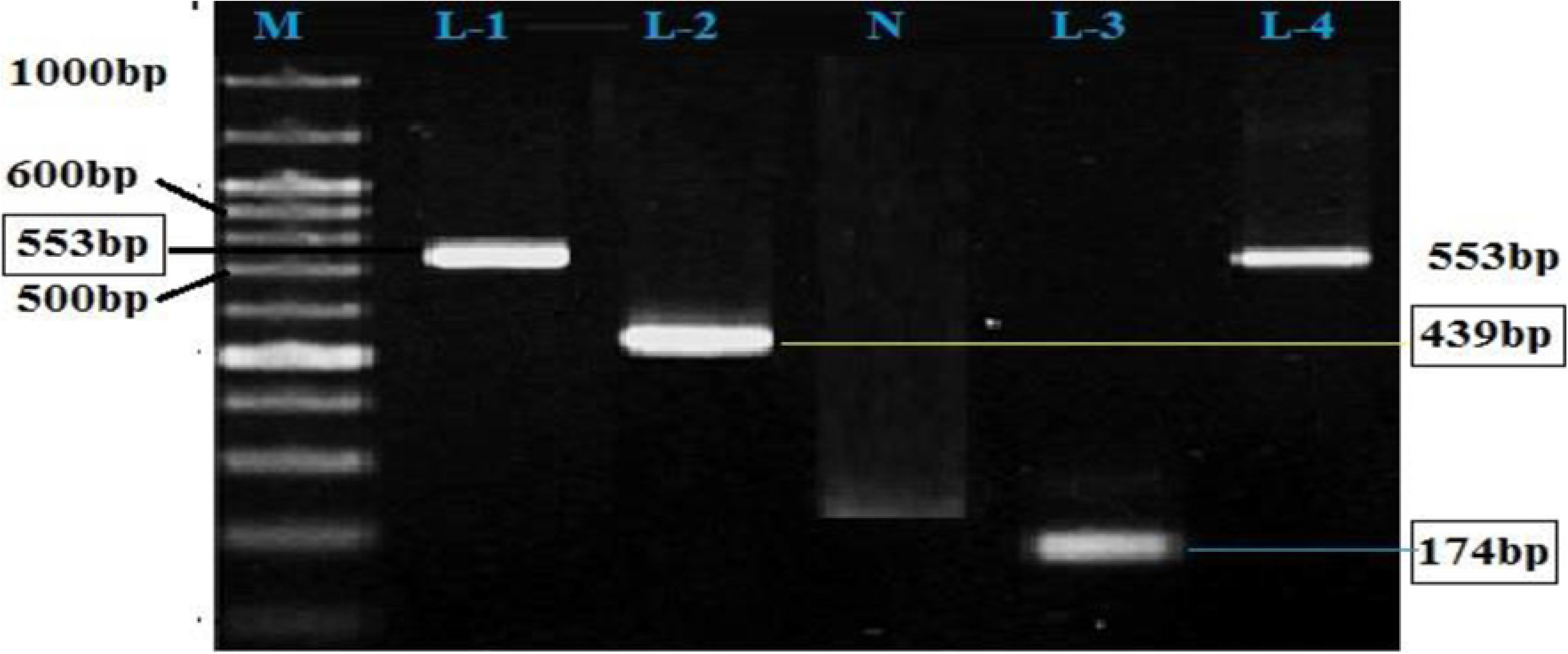
The PCR amplified product for L-1: *E. moshkovskii (*553 bp), L-2 & 4: *E. histolytica (*439 bp) and L-3: *E. dispar* (174 bp) and DNA ladder (M= marker, 1000 bp) and N= negative control.

## Discussion

The crucial importance of accurate differentiation between *E. histolytica* from *E. moshkovskii* and *E. dispar* have been reported for the actual prevalence and proper clinical management of infection in different geographical regions [25]. In different parts of Pakistan several microscopy based epidemiological surveys were performed to study the prevalence of *E. histolytica* but these studies were carried out without using molecular approaches such as PCR to accurately identify the species of *Entamoeba*. In present study a total of 400 stool samples were studied among which the 111 samples were observed positive for *Entamoeba* species. Studies on age related prevalence have been reported from Pakistan, however reports are limited showing the prevalence of *Entameoba* species among children. The overall prevalence of *Entamoeba* was recorded 27.75% (111/400). According to a previous study conducted in Multan (Punjab) Pakistan the prevalence of *Entamoeba* was recorded as 7.09% in age group 1-16 years [26]. A study from District Swat Pakistan, reported that 70 patients out 100 were infected with *E. histolytica* [27]. Another related study from Daira Ismail Khan Pakistan also reported 18.8% prevalence of *Entamoeba* species in water samples [28].

In gender wise prevalence the male participants were highly susceptible with a recorded prevalence of 33.33 % (70/210) while in female children prevalence was slightly less 21.57 % (36/190) than male. A similar study conducted in Karachi concluded that males were more affected with prevalence of 59.6% while females were found less affected with the prevalence of 40.4% [29]. The possible reason for the high prevalence in male could be the fact that male children are more exposed to environment and spent most of their time outside home which make them susceptible to infection.

The prevalence of *Entamoeba* species in children age group below 5 was 22.35 % (38/170) while in age group 5-12 year children it was observed 31.73 % (73/230). Similarly a study from Khyber Pakhtunkhwa KP, Pakistan on *E. histolytica* also reported a prevalence of 12.5% on microscopy and also concluded that the prevalence rate was higher in children below 5 years of age [30]. These findings are also an agreement with the present study from district Swat. Some related studies from Gilgit Baltistan region and Karachi city in Pakistan reported 2.5% and 8.4% of prevalence for *E. histolytica* species through microscopy [31, 32]. The prevalence rate in present study was higher due to the consumption of unhealthy and low standard food and water by the study participants. The environmental changes may also be a key factor in these findings as the weather of Gilgit is much colder than the weather of study area.

The polymerase chain reaction was used for the first time in Pakistan to differentiate the morphological same species of *Entamoeba* who shares the same morphology both trophozoit and cyst stages. The use of PCR is necessary for the differential detection of morphologically same species of *Entamoeba* because other techniques detect only *E. histolytica* while cannot differentiate among other morphological same species.

The PCR confirmed 80 out of 111 positive samples either for *E. histolytica, E. dispar* or *E. moshkovskii*. Among which the more prevalent specie was *E. dispar* in 33 (41.25%) cases followed by *E. histolytica* 25 (31.25%) and *E. moshkovskii* 7 (8.75%). The results of this research is somewhat similar to the study conducted in Iran which reported 54.8% (25/31) of all microscopic positive samples and 25 PCR confirmed cases for *E. dispar* and 8 (25.8%) positive for *E. histolytica* [33]. A study from Burkina Faso investigated 413 stool samples among which 91 were positive for *Entamoeba* species through microscopy while PCR detected only 14 samples in which the prevalence of *E. histolytica* was 21.4% (3/14) and *E. dispar* was 71.4% (10/14) [34]. A report from Malaysia shows that out of 426 stool samples only 75 were positive through microscopy while among these 75 samples 52 were observed positive through PCR in which the *E. histolytica* infection represents 75% followed by *E. dispar* 30.8% and *E. moshkovskii* 5.8% [35].

In the present study one of the important factor was the mono-infection of *E. dispar* in 33 (41.25%) cases that was higher than the pathogenic *E. histolytica* 25 (31.25%) and the co-infection of *E. dispar* with *E. histolytica* was observed in 6 (7.5%) cases. While, the co-infection with *E. moshkovskii* was 2 (2.5%) the results observed in this study shows an agreement with the study reported from India which describes that the prevalence of non-pathogenic *E. dispar* was higher 28 (23.0%) than the pathogenic *E. histolytica* 21(17.2%) while the prevalence of *E. moshkovskii* was observed in 7 (5.7%) cases [36]. A similar study was also conducted in Australia in which they subjected 110 microscopy positive samples to PCR among which 89 were amplified successfully. Out of 89 only 3 (3.4%) were found positive by PCR for *E. histolytica,* 30 (33.7%) *E. dispar* and 22 (24.7%) for *E. moshkovskii* and the co-infection of *E. dispar* with *E. moshkovskii* was 32 (26%) [37]. A study from United Arab Emirates reported that out of 120 microscopic positive samples only 23 were positive for PCR among which 12 (10%) were mono infection with *E. histolytica* 3 (2.5%), *E. dispar* and 4 (2.5%) for *E. moshkovskii* infections [38]. In present study the incidences of *E. dispar* was higher as compared to *E. histolytica* and *E. moshkovskii* therefore our findings shows a relevance with previously reported studies from other parts of the world.

Pakistan is on 3^rd^ number in world where people are suffering from GIT problems due to *E. histolytica* because the living standard and hygienic conditions of this region are very poor. Also the sanitary system is very poor and not up to the mark these may be the possible reasons for this high prevalence rate of *E. histolytica*. As we know the amoebiasis spread through contamination and *E. histolytica* is also found in animals so keeping pets and domestic animals can increase the risk of *E. histolytica* infections. It is therefore suggested that further studies are needed for identification and genetic diversity of these parasites and to determine the true pathogenicity and associated risk factors of *Entamoeba* species.

## Conclusions

In present study the PCR method was used for the first time and no such study is ever reported from Pakistan. The results of present study provides an important data for the public health care centers and clinician in Pakistan and clearly indicates the advantages of PCR over microscopy in both specificity and sensitivity. Therefore our findings suggests the use of PCR for routine-base diagnosis in this region. These results also revealed the presence of *E. dispar* and *E. moshkovskii* in district Swat and need an immediate diagnostic method in order to avoid unnecessary treatment with anti-amoebic drugs. In present study it was found that the major cause of amoebiasis in the area was poor sanitation, unhygienic conditions and consumption of junk food. Therefore, it is also suggested for awareness of local community to improve proper sanitation, hygienic conditions and healthy food.

## Acknowledgements

The authors acknowledge the support and facilitation in sampling from Al-Shifa diagnostic Lab (Saidu Sharif Swat) and Pathology Department of Saidu Medical College (SMC) Swat.

